# Differential expression of Ormdl genes in the islets of mice and humans with obesity

**DOI:** 10.1101/715599

**Authors:** Rachel Fenske, Hugo Lee, Tugce Akcan, Elliot Domask, Dawn Belt Davis, Michelle Kimple, Feyza Engin

**Author notes:** These authors contributed equally to this work (Rachel Fenske, Hugo Lee). To whom correspondence should be addressed Feyza Engin, Ph.D., Assistant Professor of Biomolecular Chemistry, HF DeLuca Biochemical Sciences Building, 440 Henry Mall, Room 6260B, Madison, WI 53706, USA.

## Abstract

The orosomucoid-like proteins (Ormdl1-3) are emerging as the critical regulators of sphingolipid homeostasis, inflammation and ER stress. However, their roles in β-cells and obesity remain unknown. Here, we showed that islets isolated from overweight/obese human donors displayed marginally reduced *ORMDL1-2* expression while *ORMDL3* expression was significantly reduced as compared to islets from lean donors. In contrast, *Ormdl3* expression was significantly upregulated in the islets of leptin-deficient obese (ob/ob) mice compared to lean mice. We identified that the difference in expression of *Ormdl3* between mouse and human islets was leptin-dependent, as treatment of *ob*/*ob* mice with leptin significantly reduced *Ormld3* expression. Furthermore, *Ormdl1-3* were significantly upregulated upon chemically-induced ER stress, but they showed differential responsiveness to cytokines in a β-cell line. Knockdown of *Ormdl3* substantially increased expression of apoptotic markers, which was rescued by a pharmacological inhibitor of ceramide synthase. Taken together we demonstrate leptin-dependent regulation of *Ormdl3* expression in ob/ob islets, highlight the possible importance of β-cell stress conditions in differential *Ormdl* expression and identify a critical role for *Ormdl3* in β-cell survival.

## Introduction

Insulin resistance, often co-incident with obesity, dampens the brake on lipolysis, elevating plasma free fatty acid levels. Free fatty acids taken up from the plasma are the precursors for various species of intracellular lipids. Certain sphingolipids, most notably, ceramide, accumulate within insulin-resistant tissues of animals^1,2^ and humans^3,4^, including the pancreatic β-cell.^5^ There, they inhibit insulin action, activating processes including apoptosis, inflammation, and stress responses—a condition known as lipotoxicity. *De novo* sphingolipid synthesis, where fatty acids from exogenous sources are utilized as substrates, is primarily responsible for obesity-induced ceramide generation^6,7^. Serine palmitoyltransferase (SPT), which catalyzes the decarboxylative condensation of L-serine and palmitoyl-CoA to 3-ketodihydrosphingosine, initiates *de novo* sphingolipid synthesis. Surplus exogenous fatty acids not only lead to increased substrate availability, but also alter expression and activity of key enzymes in the sphingolipid synthetic pathway. SPT functions as a heterodimer of subunits SPTLC1 or SPTLC2 with SPTLC3, and high fat diet feeding promotes both SPT subunit gene transcription and catalytic activity^8,9 10^. Yet, despite recent progress in the field, the molecular mechanisms of sphingolipid-mediated disease pathology and the pathways generating these pathogenic lipids remain poorly understood.

The members of the orosomucoids (Orm) gene family encode transmembrane proteins localized in the endoplasmic reticulum (ER). While the budding yeast *S. cerevisiae* has two Orm genes (Orm1 and Orm2)^11^, mammalians have three Orm-like proteins (Ormdl1-3)^12^. While Orm protein sequences are highly conserved among species, little information has been identified regarding the mechanisms of action or regulation of activity of Orm family proteins. Orm1 and Orm2 were recently identified as negative regulators of SPT. Orm proteins form a complex with SPT and inhibit its activity. This association with SPT is regulated by Orm protein phosphorylation: an important factor for sphingolipid homeostasis^13^. *In vitro* studies have suggested that mammalian Ormdl3 alters ER-mediated calcium (Ca^2+^) homeostasis, facilitates the unfolded protein response (UPR), induces cellular stress responses, and may play a role in inflammation^14-17^. In human genome wide association studies, *ORMDL3* is strongly associated with inflammatory diseases, including asthma, Crohn’s disease, and type 1 diabetes (T1D)^18-24^. Although emerging data suggest Ormdl proteins are involved in sphingolipid homeostasis, chronic inflammation, and ER stress—all of which play critical roles in the development and progression of obesity, diabetes, and β-cell dysfunction—the expression, function, and importance of Ormdl genes in β-cell physiology and pathology currently remains unknown. Here, we analyzed the expression of Ormdl genes in a β-cell line under various cellular stress conditions, in a genetic mouse model of obesity and type 2 diabetes prior to the onset of hyperglycemia, and in human pancreatic islets isolated from lean and obese non-diabetic donors. Our results for the first time, revealed obesity-, species- and sex-dependent differences in Ormdl family member expression in pancreatic islets. We also identified that expression of *Ormdl1-3* were significantly increased upon chemically-induced ER stress, but they showed differential responsiveness to cytokine-induced stress in a mouse β-cell line. Moreover, our data highlight that leptin can regulate *Ormdl3* expression and provide an explanation for differential expression of this gene in mouse and human islets in the context of obesity. Finally, we demonstrated that among the Ormdl genes knockdown of *Ormdl3* cause substantially increased β-cell death and upregulation of apoptotic markers. The cell death phenotype was markedly rescued by treatment of cells with a chemical inhibitor of ceramide synthase suggesting Ormdl3-sphingolipid-ceramide axis can play an important role in β-cell survival.

## Results

### Overweight/obese human islets display significantly reduced expression of *ORMDL3*

To identify the pancreatic islet *ORMDL* expression in the context of obesity, we used pancreatic islets isolated from lean and obese human organ donors. We grouped donors as lean (BMI≤25) and overweight/obese (BMI>25) (Table 1). All ORMDL genes showed a trend toward diminished mRNA expression in islets isolated from obese humans as compared to lean (as quantified by cycle threshold as compared to β-actin), with the cycles necessary to amplify ORMDL3 PCR product being significantly reduced (approximately 3.5 cycles, or 11-fold) (Figures 1a-c). We next examined the relationship between islet *ORMDL* expression and donor sex. Interestingly, the cycle threshold necessary to amplify *ORMDL2* and *ORMDL3* expression was significantly reduced (by approximately 5-5.5 cycles) in islets from overweight/obese female donors only, corresponding with a 32-45-fold decrease in mRNA expression with obesity (Figures 1d-f). *ORMDL1* expression level was non-significantly decreased in islets from female donors as a factor of overweight/obesity. Although no significant changes in the expression of any *ORMDL* family member were observed in islets isolated from male donors as a factor of overweight/obesity, the mean *ORMDL3* cycle threshold in islets from overweight/obese male donors was reduced as compared to lean (Figures 1d-f). To rule out a potential confounder in our human islet analyses, we examined the correlation between ORMDL expression and donor age and did not detect any significant correlation (Figures S1a-c).

**Table 1.** Description of human islet donors. Demographic and anthropometric data for each human islet donor is provided.

| **BMI Group** | **Donor** | **Age** | **Sex** | **BMI** | **Ethnicity** |
| --- | --- | --- | --- | --- | --- |
| **<25** | 1 | 40 | M | 19.6 | Black |
|  | 2 | 21 | F | 21.6 | White |
|  | 3 | 53 | M | 21.8 | White |
|  | 4 | 42 | M | 22.76 | Black |
|  | 5 | 59 | F | 23.1 | White |
|  | 6 | 51 | F | 23.1 | White |
|  | 7 | 61 | F | 23.2 | White |
|  | 8 | 48 | F | 24.19 | White |
|  | 9 | 51 | M | 24.2 | Hispanic/Latino |
| **25-29.9** | 1 | 55 | F | 25.8 | White |
|  | 2 | 36 | M | 26.0 | White |
|  | 3 | 24 | F | 26.6 | Hispanic/Latino |
|  | 4 | 60 | M | 27.3 | Asian Indian |
|  | 5 | 36 | M | 28.7 | White |
|  | 6 | 25 | M | 29.3 | Hispanic/Latino |
|  | 7 | 43 | M | 29.6 | White |
|  | 8 | 38 | M | 29.8 | Black |
| **≥30** | 1 | 29 | M | 30.2 | Asian Indian |
|  | 2 | 58 | F | 31.1 | White |
|  | 3 | 36 | M | 33.8 | White |
|  | 4 | 21 | M | 33.8 | White |
|  | 5 | 19 | M | 34.1 | White |
|  | 6 | 36 | F | 34.8 | Hispanic/Latino |
|  | 7 | 63 | M | 38.6 | White |

**Figure 1.**
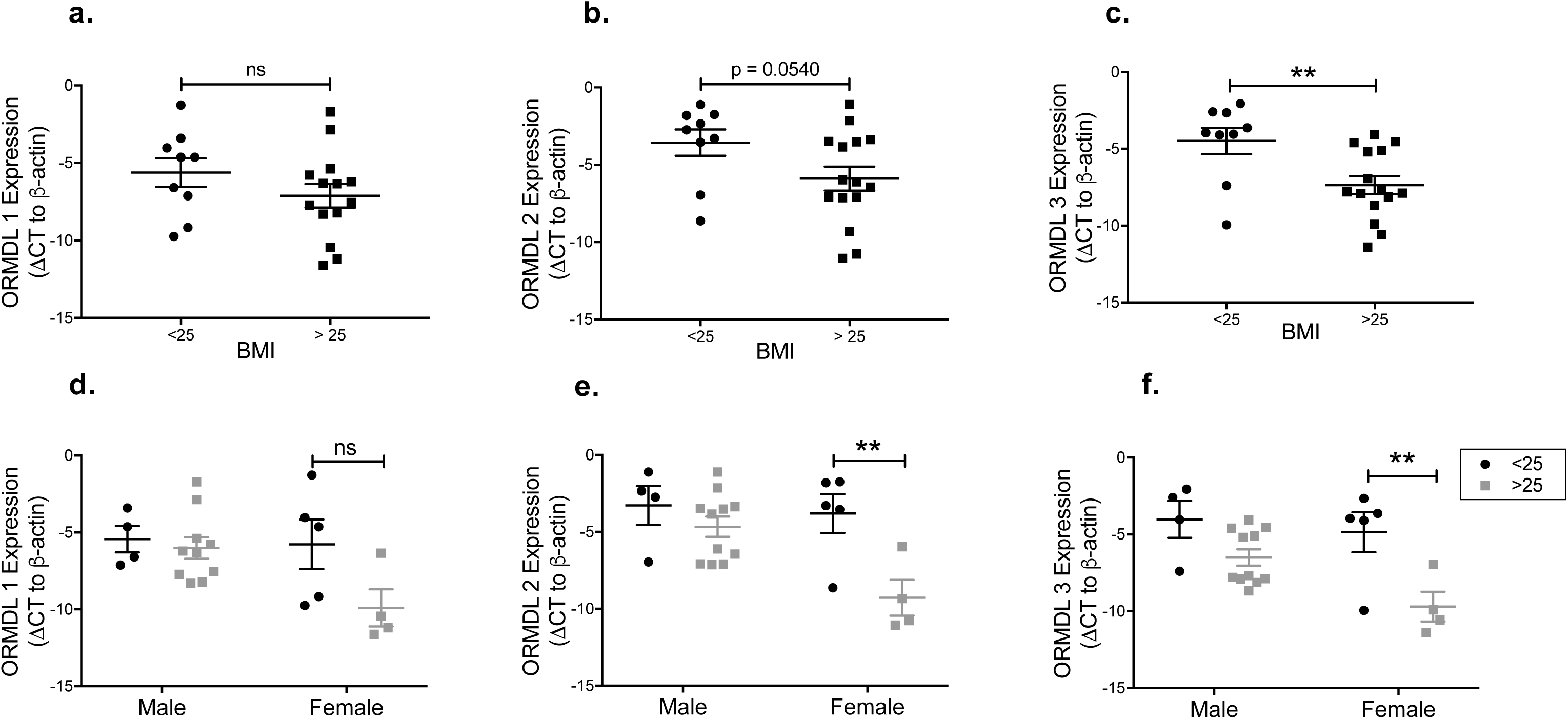
*ORMDL* gene expression in islets from overweight/obese human donors. Quantitative PCR analyses of *ORMDL1, ORMDL2*, and *ORMLD3* mRNA expression in islets from human organ donors. Expression levels of **a.** *ORMDL1* **b.** *ORMDL2* and **c.** *ORMLD3* for human donors divided into groups by BMI ≤25 (lean; n=9) (black circles) and BMI>25 (overweight/obese; n=15) (gray squares). Expression levels of **d.** *ORMDL1* **e.** *ORMDL2* and **f.** *ORMLD3* in male vs. female donors. All data are expressed as ΔCT (vs. β-actin) and are represented as mean ± SEM (\*\**p*<*0.01*). ns: non-significant.

### *Ormdl3* expression is significantly upregulated in leptin-deficient obese (ob/ob) mice

In rodent models of obesity, increased islet ceramide and triglyceride production precede β-cell dysfunction and destruction. Therefore, we examined whether the expression of Ormdl genes was altered in the pancreatic islets of genetically-induced leptin-deficient obese (ob/ob) mice, a model of severe insulin resistance and lipotoxicity. First, we analyzed the expression of Ormdl genes in lean or obese (ob/ob) male mice on C57BL/6J background at 10 weeks of age, while obese mice were still normoglycemic. Quantitative PCR analysis indicated that while expression of *Ormdl1* and *Ormdl2* non-significantly enhanced, *Ormdl3* expression was substantially upregulated in islets from male ob/ob mice as compared to lean mice (Figures 2a-c). To explore any sex-specific differences in islet *Ormdl* expression with obesity, we assessed the expression of the Ormdl genes in islets harvested from 10-week-old female C57BL/6J mice, both lean and obese. The cycles necessary to amplify *Ormdl1* and *Ormdl2* PCR product were nearly identical between islets isolated from lean and obese female C57BL/6J mice, whereas female ob/ob islets, similar to islets of male mice, showed significantly increased expression of *Ormdl3* (Figures 2d-f).

**Figure 2.**
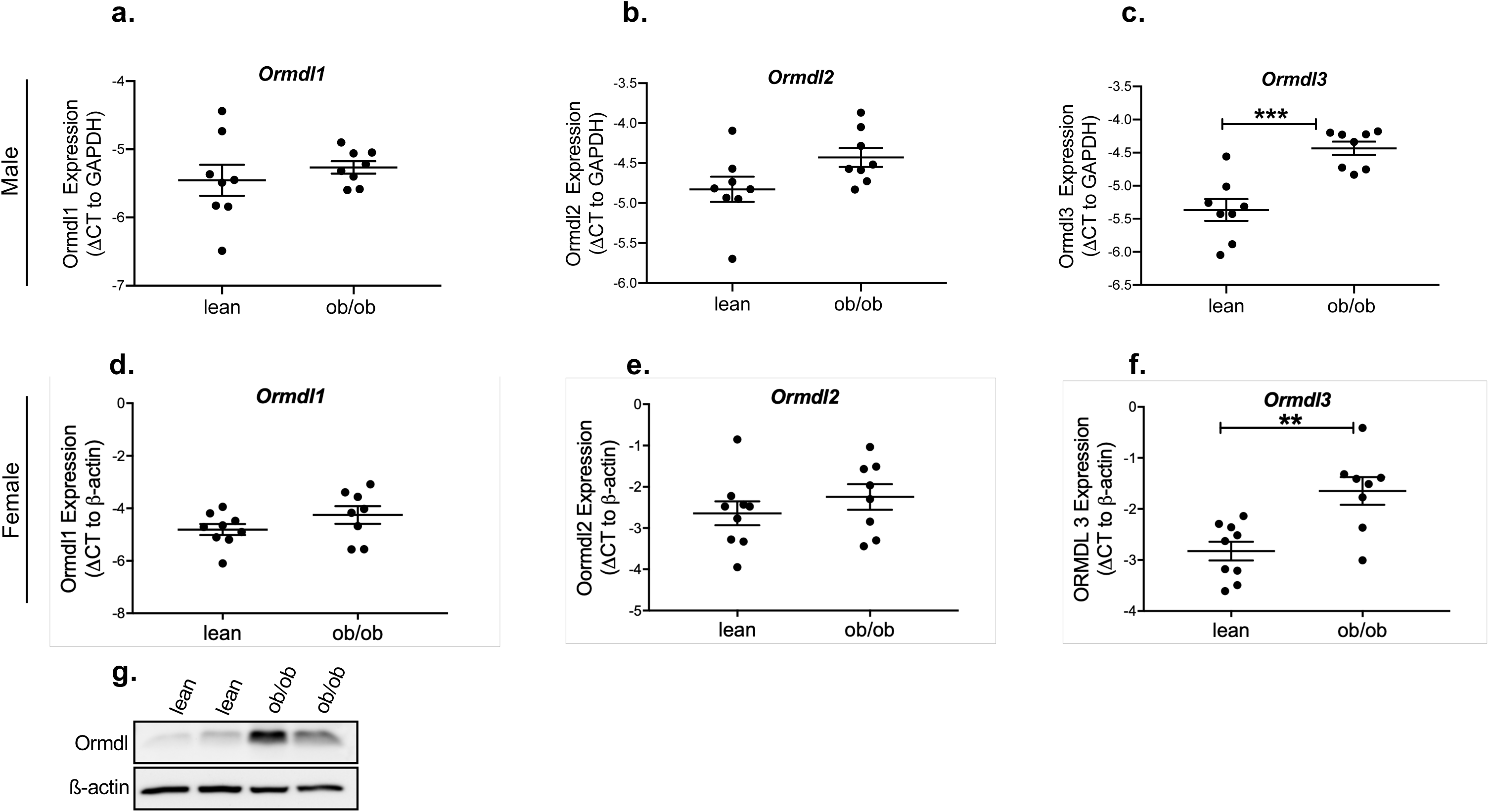
The expression of Ormdl genes in islets of leptin deficient obese (ob/ob) mice. Quantitative PCR analyses of *Ormdl1, Ormdl2* and *Ormdl3* mRNA expression in primary islets of 10-week-old male **(a-c)** and female **(d-e)** lean and obese (ob/ob) mice. All data are expressed as ΔCT (vs. β-actin or GAPDH) and represented as mean ± SEM (\*\*\**p*<*0.001*).

Next, we investigated the expression of Ormdls at the protein level. Ormdl1, −2 and −3 share greater than 80% sequence homology. Currently, there are no commercially available antibodies that can detect specific expression of the individual Ormdl family members. In addition, three commercially available pan-Ormdl antibodies failed validation using knockdown lysates (data not shown). We obtained a TPF-Ormdl antibody from Dr. Petr Draber’s group^25^ and although the antibody had significant non-specific cross-reactivity, transfection with an Ormdl3 siRNA resulted in a specific decrease in the abundance of a protein at the expected molecular weight for Ormdl (17.5 kD) that was well-removed from any other non-specific band, confirming the validity of this antibody for further analyses (Figures S2a-c). Using this antibody, we demonstrated Ormdl protein levels were also significantly upregulated in islets of male ob/ob mice, consistent with changes in mRNA expression (Figure 2g).

### Leptin administration markedly reduces *Ormdl3* expression in ob/ob islets

Our data reveal that the expression of *Ormdl3* in mouse and human islets were altered in the opposite direction. One explanation for these disparate results could have been the effect of the adipokine leptin. Obesity in humans is known to be associated with increased circulating leptin, while ob/ob mice are leptin deficient. To test this, we treated 10-week-old male normoglycemic ob/ob mice with recombinant leptin for four days. Interestingly, while we did not detect a significant effect of leptin treatment on the expression of *Ormdl1 and 2, Ormld3* expression was significantly reduced in ob/ob islets upon leptin treatment, similar to that observed in human islets from obese donors as compared to lean (Figures 3a-c). Mean blood glucose levels in ob/ob mice were not significantly changed by leptin treatment, but an approximate 10% reduction in bodyweight was observed, as previously reported^26,27^ (Figures 3d and 3e). These results indicate that the leptin can play a key role in *Ormdl3* transcriptional regulation and provide an explanation for the differential expression pattern that we observed in islets obtained from overweight/obese human donors and leptin deficient ob/ob mice.

**Figure 3.**
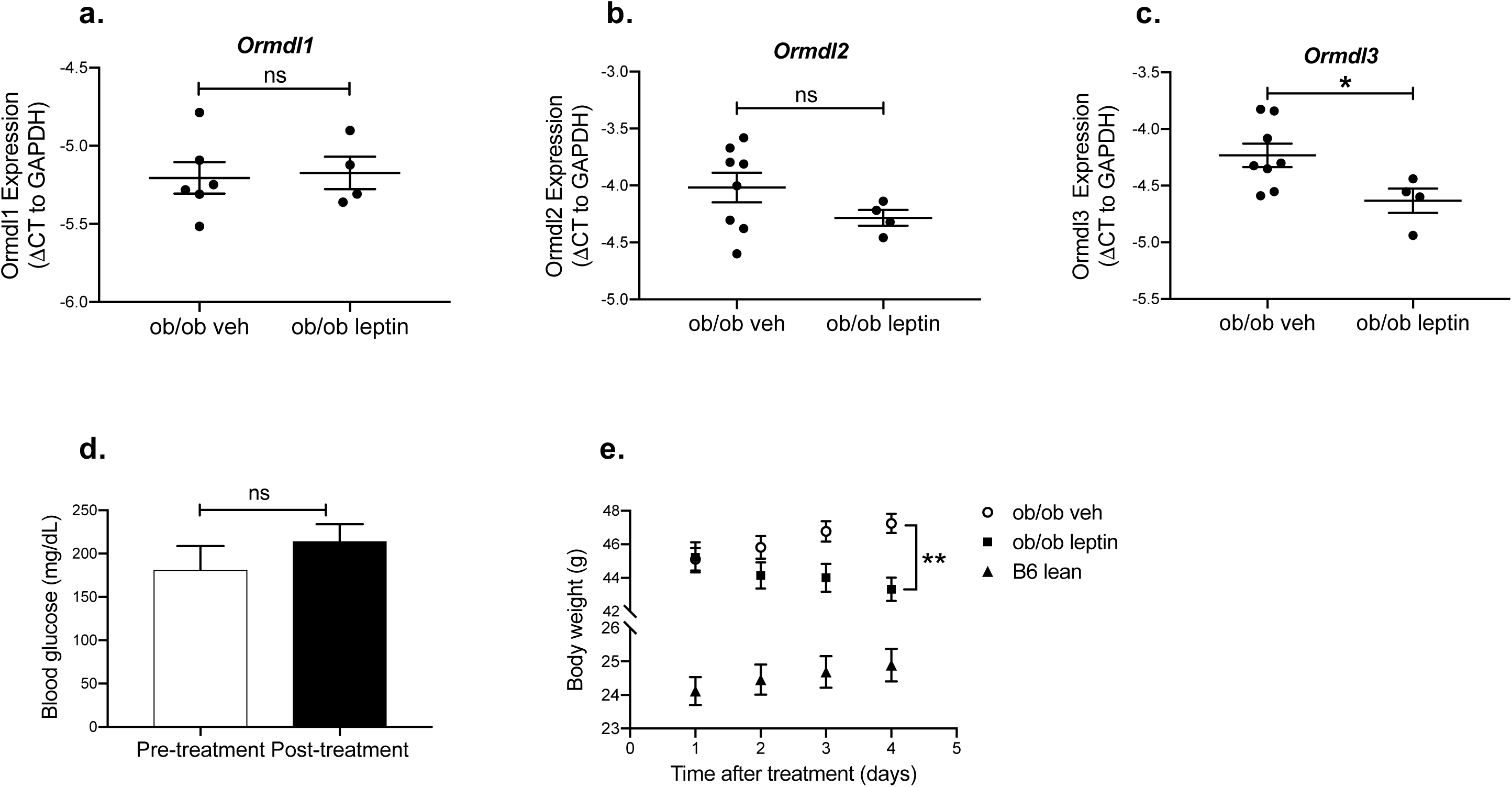
Treatment of ob/ob mice with recombinant leptin. Quantitative PCR analyses of **a.** *Ormdl1*, **b.** *Ormdl2*, and **c.** *Ormdl3* mRNA expression in primary islets of 10-week-old male obese (ob/ob) mice treated with leptin (n=4) or vehicle (n=8). **d.** Fasting blood glucose levels of ob/ob mice before and after leptin treatment. **e.** Daily body weight measurements of ob/ob and age, sex-matched control C57BL/6J mice (n=12). All data are expressed as ΔCT (vs. GAPDH) and represented as mean ± SEM (*p<0.05, **p<0.01). ns: non-significant.

### Chemically-induced ER stress upregulates the expression of *Ormdl1-3*

Accumulation of the sphingolipid, ceramide, can lead to ER stress. Localized ceramide production and an associated decrease in ER sphingomyelin levels in response to lipotoxicity is an important initiator of ER stress in the β-cell. To evaluate whether ER stress itself can alter β-cell expression of Ormdl genes, we first treated a mouse insulinoma cell line, MIN6, with a chemical ER stress inducer, tunicamycin, for 24 hours. Tunicamycin-treated MIN6 cells had higher expression of all three Ormdl family members than control-treated cells, with expression of Ormdl1, 2 and Ormdl3 being significantly increased (by approximately 3-fold) (Figures 4a-c). In contrast, treating MIN6 cells with a pro-inflammatory cytokine cocktail had little-to-no impact on Ormdl mRNA expression, save a small (1.2-fold) but statistically significant increase in Ormdl1 mRNA (Figures 4a-c). Next, we examined the effects of thapsigargin, cytokines, and an additional ER stressor, tunicamycin, on total Ormdl protein expression at 8- and 24 hours post-treatment in MIN6 cells. We confirmed thapsigargin and tunicamycin upregulated the ER stress pathway by Western blot for spliced Xbp1 (sXbp1) and the chaperone, GRP78. sXbp1 was increased at 8 hours and GRP78 at 24 hours with thapsigargin and tunicamycin treatment, as expected (Figures 4d and 4e). Total Ormdl expression was significantly increased 8 hours after initiation of treatment with ER stressors (Figure 4d), and, at 24 hours, Ormdl protein expression was markedly increased upon induction of ER stress (Figure 4e). Despite these changes in expression levels with chemical ER stress inducers, we have not detected any significant alterations in the protein levels of Ormdl genes at the indicated time points upon cytokine cocktail treatment.

**Figure 4.**
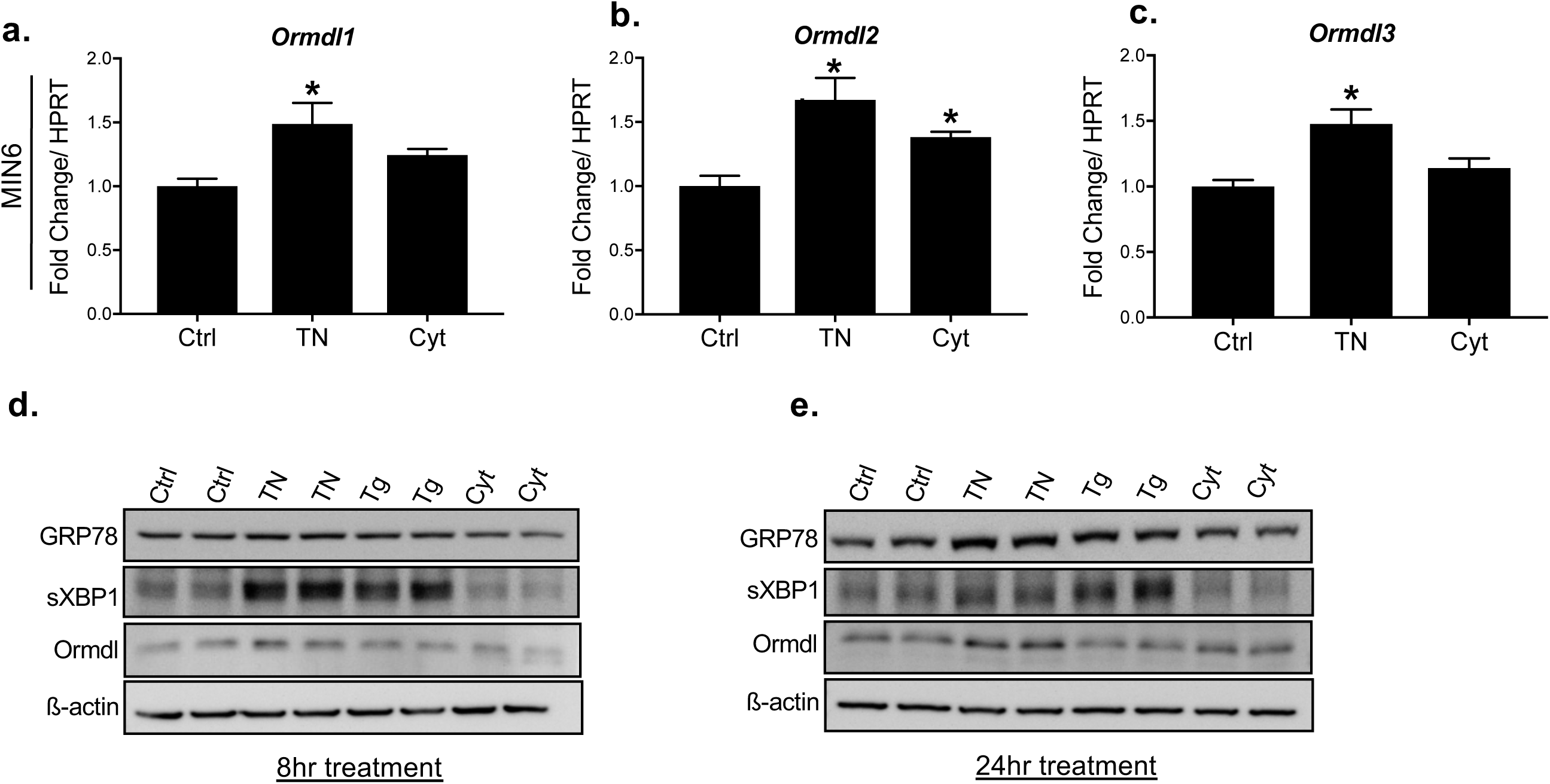
Time course analysis of total Ormdl protein levels in a mouse β-cell line upon induction of cellular stress. MIN6 cells in triplicate wells were treated with DMSO (vehicle control) or 1µM tunicamycin (TN) or a cytokine cocktail (TNF-α, IL-1β, and IFNγ) for 24 hours. Following RNA extraction, changes in **a.** *Ormdl1*, **b.** *Ormdl2* and **c.** *Ormdl3* mRNA levels were determined by qPCR. MIN6 cells were treated with tunicamycin (TN), thapsigargin (Tg) and a cytokine cocktail for **d.** 8 hours and **e.** 24 hours. Grp78, spliced Xbp1 (sXbp1), total Ormdl, and beta-actin protein levels were determined by Western blot. All data are expressed as fold change relative to control treatment, and are represented as mean ± SEM (\**p*<*0.05*). Ctrl: Control, TN: Tunicamycin, Cyt: Cytokine cocktail.

### Knockdown of *Ormdl3* leads to β-cell death

Next, to investigate the physiological function of Ormdl genes in β-cells we knocked down all three members of the *Ormdl* family in the INS1-derived 832/3 rat insulinoma cell line using siRNAs specific to each gene product. A knockdown efficiency of 80-85% was confirmed by qPCR for each of the Ormdl genes (Figures 5a-c). 48 hours after knockdown, more than half of the *Ormdl3*-deficient cells were visibly dead with appearance of floating cells. We assessed expression levels of apoptosis markers in these cells by Western blot. Apoptotic markers caspase-3 and cleaved Parp were moderately increased in *Ormdl2* deficient cells, while *Ormdl1*-deficient cells appeared to have only marginally increased cleaved Parp expression (Figures 5d and 5e). Consistent with our observations of floating dead cells, we identified significantly increased levels of caspase-3 and cleaved Parp in *Ormdl3* knockdown cells (Figure 5f). These data suggested that Ormdl3 can play a significant role in β-cell survival under physiological conditions.

**Figure 5:**
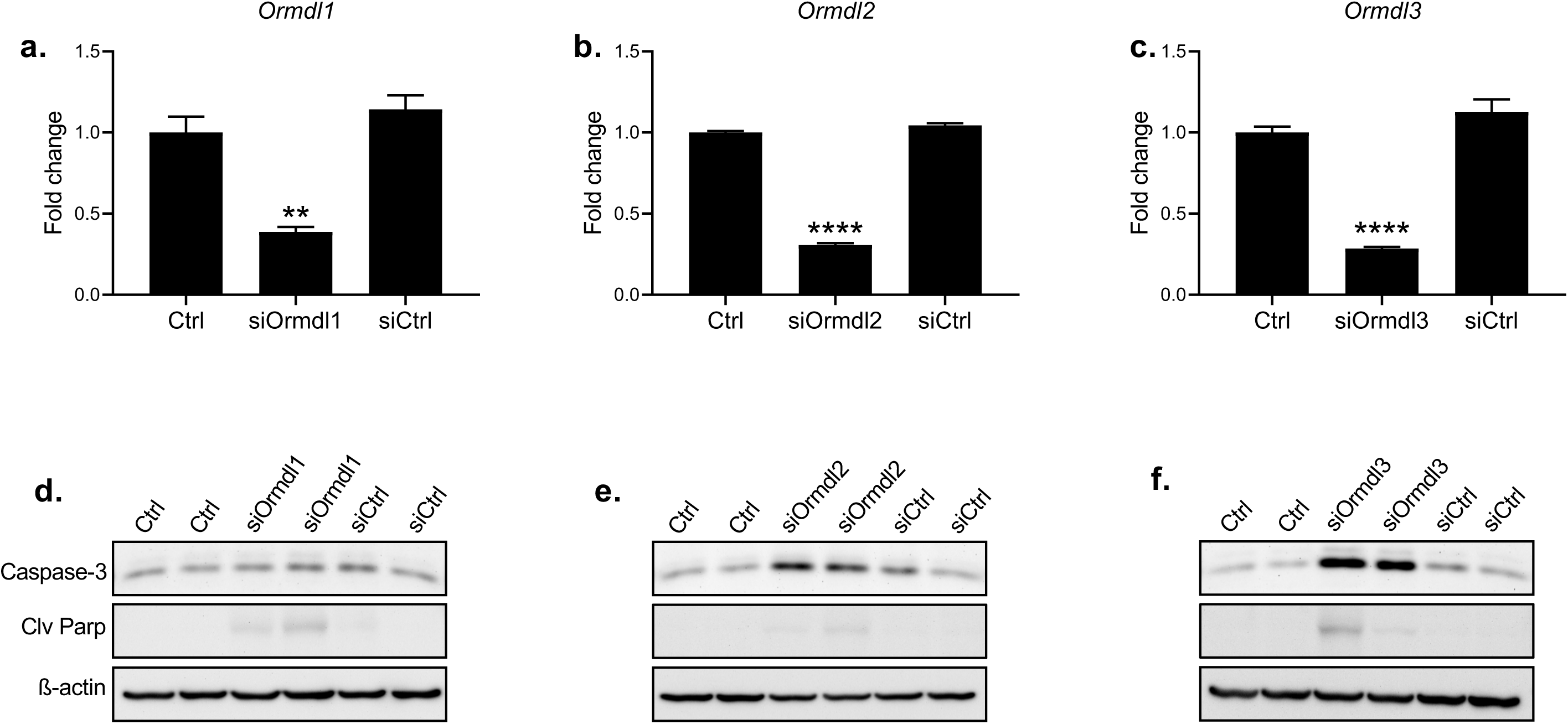
Knockdown of Ormdl genes in a β-cell line. INS-1 832/3 β-cells in triplicate wells were transfected with **a.** *Ormdl1*, **b.** *Ormd2* and **c.** *Ormdl3* siRNAs and the efficiency of knockdowns were determined by qPCR 24 hours after the transfection. The expression levels of apoptotic markers, caspase-3 and cleaved Parp in **d.** *Ormdl1*-, **e.** *Ormdl2*- and **f.** *Ormdl3*- deficient cells were assessed by Western blotting. Representative blots from two biological replicates are included from a total of three different experiments. All data are expressed as fold changes relative to baseline and represented as mean ± SEM, with statistical analysis performed by Student’s t test (\*\*\*\**p*<*0.0001*, \*\**p*<*0.01*).

### β-cell death induced by *Ormdl3* deficiency do not alter the UPR, but can be rescued by a ceramide synthase inhibitor

Ormdl3 has been reported to regulate ER stress and the UPR. Thus, we investigated whether β-cell death observed in Ormdl3 deficient cells was due to increased ER stress and/or dysfunctional UPR. We did not observe any significant changes in the mRNA or protein levels of the UPR markers sXbp1, Grp78, Chop or Atf6 in INS-1 832/3 cells transfected with siOrmdl3 alone or in the presence of ER stressor thapsigargin (Figures 6a-e). These data suggest that Ormdl3 deficiency does not trigger ER stress or ER stress-mediated cell death in INS-1 832/3 cells. Next, we hypothesized that Ormdl3 deficiency might increase ceramide production and subsequently lead to β-cell death as Orm proteins known to suppress sphingolipid biosynthesis and ceramides, which are known to be important mediators β-cell dysfunction and apoptosis^21,22^. If this is the case, blocking the ceramide synthesis downstream of sphingolipid synthesis by using a pharmacological inhibitor of ceramide synthase, Fumonisin B1, should have reduced the cell death. To test this hypothesis, we treated Ormdl3 deficient INS-1 cells with Fumonisin B1, an inhibitor of *de novo* ceramide biosynthesis, and assessed protein levels of apoptosis markers via western blotting. Consistent with our hypothesis, levels apoptotic markers, caspase-3 and cleaved-Parp, were markedly reduced upon inhibition of ceramide synthase in INS-1 832/3 cells (Figure 6f).

**Figure 6:**
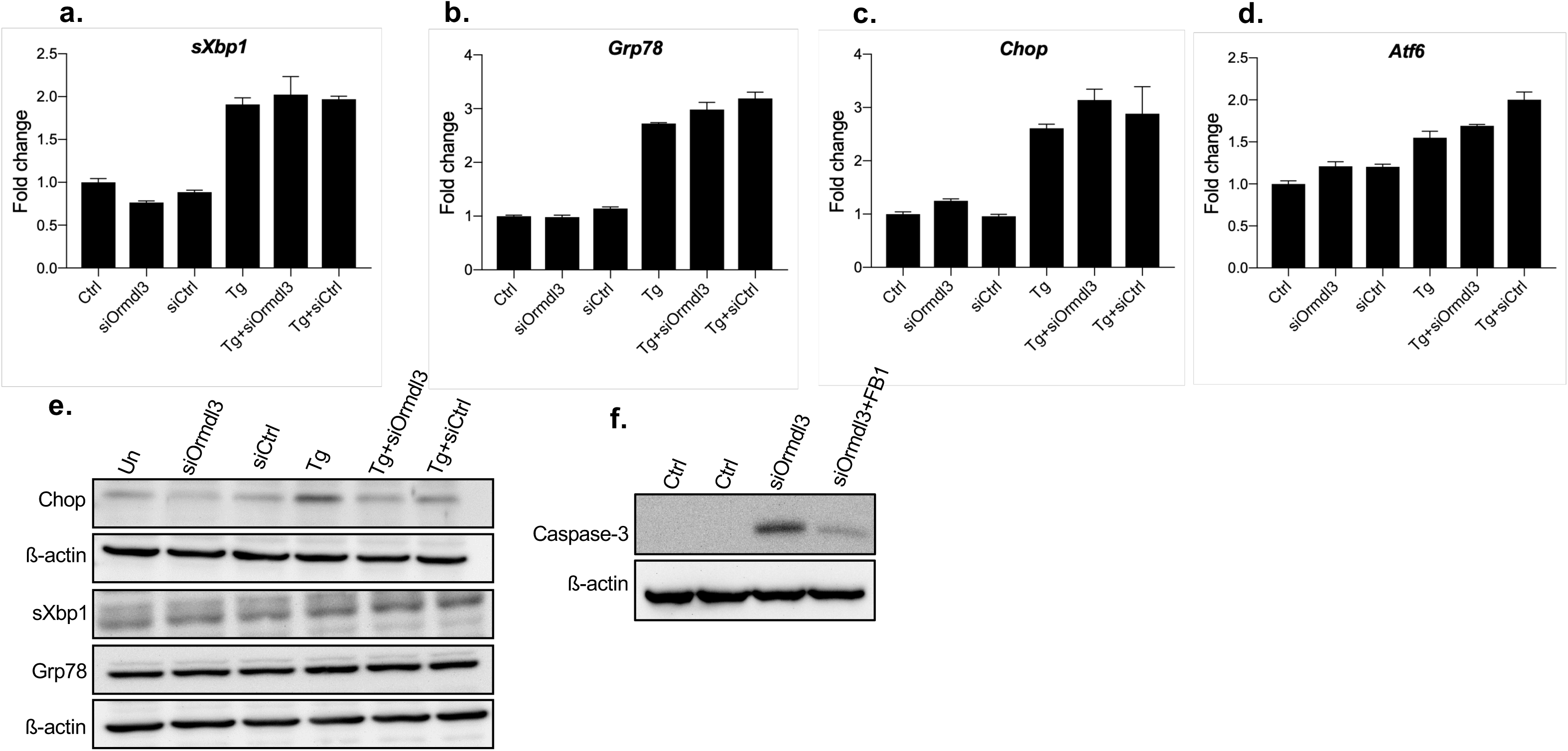
The effects of Ormdl3 knockdown on ER stress. INS-1 832/3 cells were transfected with siOrmdl3 and 24 hours after transfection cells were treated with 10nM Tg for 8 hrs. RNA was collected and qPCR was performed for **a.** sXbp1, **b.** Grp78 **c.** Chop, **d.** Atf6. **e.** Cells were transfected with siOrmdl3 and protein expression of Chop, sXbp1 and Grp78 was determined by western blotting. **f.** INS-1 832/3 cells transfected with siOrmdl3 and treated with 15µM Fumonisin B1 for 24hrs. Apoptotic markers were assessed by examining the protein expression of caspase-3 and cleaved-Parp by western blotting.

## Discussion

A growing body of evidence has suggested the genetic association of ORMDL3 gene polymorphisms with a diverse set of inflammatory disorders that include bronchial asthma, inflammatory bowel disease, ankylosing spondylitis, T1D, and atherosclerosis. We have recently shown aberrant β-cell ER stress is linked to T1D pathogenesis, and that there is abnormal β-cell UPR activity in both type 1 and type 2 diabetes animal models and human patients suggesting that highly conserved cellular mechanisms can play a critical role in the pathology of both types of diabetes^28-30^. Hence, due to their involvement with inflammatory diseases, regulation of sphingolipid biosynthesis, and ER stress we hypothesized that Ormdls could play an important role in β-cell homeostasis, and we investigated the regulation of these genes in pancreatic islets in the context of obesity. One of the most intriguing findings of our study was islet *ORMDL3* expression was significantly influenced by obesity in both mouse and human samples, albeit in opposing directions. *ORMDL3* mRNA expression was significantly reduced in islets isolated from overweight/obese human female organ donors while *Ormdl3* expression was actually increased in islets from both female and male *ob*/*ob* mice, the latter significantly. We reasoned that these contrasting results might be due to leptin as while obese humans have significantly increased levels of circulating leptin, ob/ob mice are deficient of this adipokine. Indeed, administration of leptin to male ob/ob mice only for four days significantly reduced *Ormdl3* expression in islets, indicating that leptin may have a regulatory role in Ormdl3 expression. Interestingly, central leptin signaling represses SPT expression in white adipose tissue by 30%^31^ and decreases mRNA expression of enzymes involved in *de novo* ceramide synthesis (SPT-1, LASS2, LASS4) and ceramide production from sphingomyelin (SMPD-1/2)^31^. Leptin receptor overexpression in the islets of obese Zucker diabetic fatty (ZDF) rats with mutant leptin receptors leads to significantly reduced SPT mRNA and fat content^32,33^. However, whether such a regulation also exists in human islets is not yet known. Since ob/ob mice have elevated circulating free fatty acids and increased ceramide in their islets^34^, it is possible that upregulated *Ormdl3* levels in ob/ob mice in the absence of leptin exert compensatory inhibitory effects on sphingolipid synthesis, whereas in obese, hyperleptinemic human subjects, downregulation of *ORMDL3* may lead to increased ceramide synthesis and lipotoxicity in islets. Of note, while leptin resistance in the hypothalamus is well established in obesity and diabetes, such resistance has not been definitively demonstrated in pancreatic β-cells. Leptin levels are also higher in women than in men using any given measure of obesity; a finding that may provide insight into the sex-dependent differential expression of *ORMLD2* and *ORMDL3*.

Recent studies suggested the genetic association of *ORMDL3* polymorphisms with a diverse set of inflammatory disorders that include bronchial asthma, inflammatory bowel disease, ankylosing spondylitis, T1D, and atherosclerosis. Hence, we tested whether cytokines and ER stress, which play a key role in the pathogenesis of many inflammatory diseases would affect Ormdl expression. Interestingly, while we did not detect a major change in the expression of Ormdl genes in MIN6 cells in response to cytokines, however chemically-induced ER stress significantly upregulated *Ormdl3* in these cells. Since cytokines known to induce ER stress, whether under prolonged cytokine exposure and various dose regimes inflammatory cytokines alter Ormdl gene expression in an ER stress-dependent matter will need further investigation. Emerging data indicate that Ormdl3 promotes ER stress and the UPR^15,16^, here we show that *Ormdl3* itself can be regulated by chemically-induced ER stress, but not induce ER stress in a β-cell line. In addition, we demonstrated that even in the absence of any additional stressor, deficiency of *Ormdl3* leads to β-cell death and upregulation of apoptotic markers in INS-1 832/3 cells which can be significantly reduced by administration of a pharmacological inhibitor of ceramide synthesis. This demonstrates for the first time that Ormdl3 plays an important role in β-cell survival. However, the exact function and regulation of Ormdl3 in β-cells will need to be identified using tissue-specific genetic loss and gain of function models.

Our findings do not clarify whether Ormdl protein function itself was altered under the conditions of obesity and/or inflammation. In yeast, Orm proteins are regulated through phosphorylation^35,36^. Although, Orm proteins are highly conserved in higher organisms, Ormdl proteins have a truncated N-terminal domain and lack the phosphorylation motif found in yeast, indicating divergent post-translational regulatory mechanisms^37^. Emerging data indicate that Ormdl genes might be transcriptionally regulated: when sphingolipid degradation was compromised by deletion of the lyase enzyme, low SPT activity paralleled a considerable elevation in Ormdl1 and Ormdl3 transcription^38^. In addition, ORMDL expression strongly influenced SPT activity and *de novo* ceramide synthesis in macrophages^39^. The molecular mechanisms of regulation of Ormdl transcription and post-translational processing still needs detailed investigation.

To summarize, our data provide the first comprehensive analysis of *ORMDL* family gene expression in mouse and human islets in the context of obesity. Furthermore, we demonstrate that *Ormdl* gene expression is responsive to ER stress, a cellular stress condition that is associated with obesity, in a mouse β-cell line. We also demonstrate that leptin can have a significant role in the regulation of islet expression of *Ormdl3*. Finally, our results indicate that loss of *Ormdl3* can lead to significantly increased β-cell death in a rat insulinoma cell line possibly due to increased ceramide synthesis. The molecular mechanisms by which Ormdl proteins regulate β-cell homeostasis, under physiological and pathological conditions including obesity and diabetes remain to be determined and are worthy of future study.

### Limitations of the study

While our study reveals for the first time the expression of the Ormdl genes in the islets of both obese mice and humans, it does not address the protein levels in human samples mainly due to the limited sources of human samples and the specific antibody. Further mechanistic studies by using primary islets and *in vivo* loss and gain of function models will be necessary to definitively demonstrate the function and regulation of these genes in β-cell pathophysiology.

## Methods

### Mice

9 weeks of age C57BL/6J and ob/ob (B6.Cg-*Lep*^*ob*^/J) mice were purchased from the Jackson Laboratory and were housed under standard conditions, under a 12:12-hr light/dark cycle, with unrestricted access to food and drinking water in an animal housing facility accredited by the Association for Assessment and Accreditation of Laboratory Animal Care. This study was carried out in accordance with the recommendations of the National Institutes of Health Guide for the Care and Use of Laboratory Animals. The protocol (#M005064-R01-A03 by F.E. for mice) was approved by the University of Wisconsin-Madison Institutional Animal Care and Use Committee.

### Leptin administration

B6.Cg-Lep^ob^/J (ob/ob) male mice were purchased from Jackson Laboratories at 9 weeks of age and were assigned to either a vehicle (n=8) or leptin treatment (n=4) group. Prior and post treatment 6-hour fasting blood glucose was measured using a Breeze2 glucometer (Bayer). At 10 weeks of age, recombinant murine leptin (PeproTech, Rocky Hill, NJ, USA) was reconstituted according to manufacturer’s instructions and IP injected once daily to mice in the leptin group at a concentration of 4.5 µg/g body weight for 4 days, while mice in the vehicle group received filter-sterilized water. The body weight of vehicle and leptin-treated ob/ob mice as well as was control age, sex matched C57BL/6J mice (n=12) measured daily.

### Cell culture

MIN6 cells were cultured in DMEM supplemented with penicillin, streptomycin, 1mM sodium pyruvate, 50 μM β-ME, and 15% FBS. INS-1 832/3 cells were cultured in RPMI 1640 supplemented with penicillin, streptomycin, 2 mM glutamine, 10mM HEPES, 1mM sodium pyruvate, 50 μM β-ME, and 10% FBS. The cells were maintained at 37°C in 5% CO_2_ atmosphere and treated with 2 µg/ml tunicamycin (Sigma-Aldrich), 10 nM-1µM thapsigargin (Sigma-Aldrich), and the cytokines for MIN6: TNF-α (200 U/mL), IL-1β (20 U/mL), and IFNγ (200 U/mL), (PeproTech, Rocky Hill, NJ, USA). Experiments were carried out with the approval of the University of Wisconsin-Madison Institutional Biosafety Committee (#B00000036). For gene-specific, siRNA-mediated knockdowns, 1×10^6^ cells/well were used to perform reverse transfections. Transient transfections were carried out with 100 nM siRNA oligonucleotide pools (Sigma-Aldrich) using HiPerFect transfection reagent (Qiagen) per manufacturer’s recommendations. Transfections were carried out for 24-48 hours depending on the subsequent treatment protocol. siRNA transfections resulted in between 75–90% depletion of Ormdl mRNAs as measured by qPCR. The knockdown experiments were repeated more than three times.

### RNA extraction and qPCR analysis

Total RNA was extracted using TRIzol reagent (Invitrogen) from MIN6 cells. cDNAs were synthesized from extracted RNA by using Superscript III First Strand RT-PCR kit (Invitrogen). Real-time quantitative PCR amplifications were performed on CFX96 Touch Real-time PCR detection system (Bio-Rad). β-actin, HPRT and GAPDH genes were used as internal controls for the quantity and quality of the cDNAs in real time PCR assays. Primer specific for mouse: m*Ormdl1:* F: ACA GTG AGG TAA ACC CCA ATA CT, R: GCA AAA ACA CAT ACA TCC CCA GA; m*Ormdl2:* F: CAC AGC GAA GTA AAC CCC AAC, R: AGG GTC CAG ACA ACA GGA ATG: *mOrmdl3:* F: CCA ACC TTA TCC ACA ACC TGG, R: GAC CCC GTA GTC CAT CTG C. Primers specific for rat: r*Ormdl1* F: CCC AAT ACT CGT GTA ATG AAT AGC, R: GGG ATG TG AGA AAT ACA ATG TG; r*Ormdl2:* F: GAT GGA CTA CGG ACT ACA GTT TAC, R: AGT GAG GCA GTG TTG ATG AG; r*Ormdl3:* F: TTG ACC ATC ACG CCC ATT, R: AGC ACA CTC ATC AAG GAC AC; r*sXbp1*: F: CTG AGT CCG AAT CAG GTG CAG, R: ATC CAT GGG AAG ATG TTC TGG; *rGrp78*: F: TGG GTA CAT TTG ATC TGA CTG GA, R: CTC AAA GGT GAC TTC AAT CTG GG; *rChop*: F: CCA GCA GAG GTC ACA AGC AC, R: CGC ACT GAC CAC TCT GTT TC; *rAtf6*: F: TCG CCT TTT AGT CCG GTT CTT, R: GGC TCC ATA TGT CTG ACT CC.

Human islet RNA was extracted using the Qiagen RNeasy Kit (Qiagen; #74106) according to manufacturer’s instructions. cDNA was generated with random hexamers (High-Capacity cDNA Reverse Transcription Kit, Applied Biosystems; #4368813). qPCR was performed using SYBR green (Roche; #04913914001). Primer specific for humans: h*ORMDL1:* F: TGA CCA GGG TAA AGC AAG GC, R: CCG AAC ACC ATG TAG TTG TGG; h*ORMDL2:* F: GTG GCA CAC AGC GAA GTA AAC, R: TGC AGC AAT CCT ACC AAG ATG; h*ORMDL3*: F: GAG GCT GCT AAC CCA CTG G, R: GGT GAG GAA GTA CAG CAC GAT. All human islet cycle thresholds were normalized to β-actin.

### Donor human islets

Human islets were obtained from the Integrated Islet Distribution Program (IIDP) according to an approved IRB exemption protocol stating this work is not human subjects research (UW 2012–0865). Islets were cultured in RPMI 1640 with 8 mM glucose for 24 hours before being pelleted for RNA.

### Primary mouse islet Isolation

Islets were isolated by using the standard collagenase/protease digestion method. Briefly, the pancreatic duct was cannulated and distended with collagenase solution using Collagenase type XI (Sigma-Aldrich). Islets were separated from the exocrine tissue using Histopaque (Sigma-Aldrich) gradient. Hand-picked islets were counted prior to the experiments.

### Western Blot

Cells or islets were lysed in RIPA buffer (50 mM Tris, pH 7.4, 150 mM, NaCl, 5 mM EDTA, pH 8.0, 30 mM NaF, 1 mM Na3VO4, 40 mM β-glycerophosphate, 0.1 mM PMSF, protease inhibitors, 10% glycerol and 1% Nonidet-P40). The concentration of the isolated proteins was determined using BCA Protein Assay Reagent (Pierce, Rockford, IL). Thirty to forty-five micrograms of the protein were separated on a 5-12% Tris-acetate gel and electrophoretically transferred to PVDF membranes (Millipore, Billerica, MA). Membranes where then incubated with the primary antibodies against sXbp1 (BioLegend), Chop (Santa Cruz Biotechnology), GRP78 (Cell Signaling Technology), Caspase-3 (Cell Signaling Technology), Cleaved-PARP (Cell Signaling Technology), Ormdl (TPF, gift of Dr. Petr Draber, Academy of Sciences of the Czech Republic), β-actin (Cell Signaling Technology) and the appropriate secondary antibodies.

### Statistical analysis

Data analysis was performed using GraphPad Prism v.8 (GraphPad Software, San Diego, CA). Following Shapiro-Wilks normality testing, data were analyzed by Student’s *t*-test, unless otherwise stated. *P* < 0.05 was considered statistically significant. Data are represented as mean ± SEM or SD.

## Acknowledgements

We thank Dr. Petr Draber for providing the Ormdl antibody, and Mieke Baan for her technical help. This work was supported by grants from the JDRF-5-CDA-2014-184-A-N and NIH 5K01DK102488-03 (to F.E); NIH R01 DK102598 and VA BLR&D I01 BX003700 (to M.E.K); VA BLR&D I01 BX001880 and NIH R01 DK110324 (to D.B.D). RJF is supported by NIH F31 DK109698. H.L is supported by NIH National Research Service Award T32 GM007215. This work was supported using facilities and resources from the William S. Middleton Memorial Veterans Hospital, and does not represent the views of the Department of Veterans Affairs or the United States Government.

## Author Contribution

The hypothesis and research concept were by F.E. F.E. supervised research, analyzed the data and wrote the article. R.F., H.L., T.A., and E.D. contributed to experiments. M.E.K., D.B.D. analyzed the data and supervised research. M.E.K, D.B.D. and F.E edited the final version of the manuscript.

## References

1 Holland, W. L. & Summers, S. A. Sphingolipids, insulin resistance, and metabolic disease: new insights from in vivo manipulation of sphingolipid metabolism. Endocr Rev 29, 381–402, doi:10.1210/er.2007-0025 (2008).

2 Summers, S. A. Ceramides in insulin resistance and lipotoxicity. Prog Lipid Res 45, 42–72, doi:10.1016/j.plipres.2005.11.002 (2006).

3 Adams, J. M., 2nd et al. Ceramide content is increased in skeletal muscle from obese insulin-resistant humans. Diabetes 53, 25–31 (2004).

4 Straczkowski, M. et al. Increased skeletal muscle ceramide level in men at risk of developing type 2 diabetes. Diabetologia 50, 2366–2373, doi:10.1007/s00125-007-0781-2 (2007).

5 DeFronzo, R. A. Dysfunctional fat cells, lipotoxicity and type 2 diabetes. Int J Clin Pract Suppl, 9–21 (2004).

6 Hu, W., Bielawski, J., Samad, F., Merrill, A. H., Jr. & Cowart, L. A. Palmitate increases sphingosine-1-phosphate in C2C12 myotubes via upregulation of sphingosine kinase message and activity. J Lipid Res 50, 1852–1862, doi:10.1194/jlr.M800635-JLR200 (2009).

7 Watt, M. J. et al. Regulation of plasma ceramide levels with fatty acid oversupply: evidence that the liver detects and secretes de novo synthesised ceramide. Diabetologia 55, 2741–2746, doi:10.1007/s00125-012-2649-3 (2012).

8 Cinar, R. et al. Hepatic cannabinoid-1 receptors mediate diet-induced insulin resistance by increasing de novo synthesis of long-chain ceramides. Hepatology 59, 143–153, doi:10.1002/hep.26606 (2014).

9 Longato, L., Tong, M., Wands, J. R. & de la Monte, S. M. High fat diet induced hepatic steatosis and insulin resistance: Role of dysregulated ceramide metabolism. Hepatol Res 42, 412–427, doi:10.1111/j.1872-034X.2011.00934.x (2012).

10 Blachnio-Zabielska, A., Baranowski, M., Zabielski, P. & Gorski, J. Effect of high fat diet enriched with unsaturated and diet rich in saturated fatty acids on sphingolipid metabolism in rat skeletal muscle. J Cell Physiol 225, 786–791, doi:10.1002/jcp.22283 (2010).

11 Han, S., Lone, M. A., Schneiter, R. & Chang, A. Orm1 and Orm2 are conserved endoplasmic reticulum membrane proteins regulating lipid homeostasis and protein quality control. Proc Natl Acad Sci U S A 107, 5851–5856, doi:10.1073/pnas.0911617107 (2010).

12 Hjelmqvist, L. et al. ORMDL proteins are a conserved new family of endoplasmic reticulum membrane proteins. Genome Biol 3, RESEARCH0027 (2002).

13 Breslow, D. K. et al. Orm family proteins mediate sphingolipid homeostasis. Nature 463, 1048–1053, doi:10.1038/nature08787 (2010).

14 Miller, M. et al. ORMDL3 is an inducible lung epithelial gene regulating metalloproteases, chemokines, OAS, and ATF6. Proc Natl Acad Sci U S A 109, 16648–16653, doi:10.1073/pnas.1204151109 (2012).

15 Hsu, K. J. & Turvey, S. E. Functional analysis of the impact of ORMDL3 expression on inflammation and activation of the unfolded protein response in human airway epithelial cells. Allergy Asthma Clin Immunol 9, 4, doi:10.1186/1710-1492-9-4 (2013).

16 Cantero-Recasens, G., Fandos, C., Rubio-Moscardo, F., Valverde, M. A. & Vicente, R. The asthma-associated ORMDL3 gene product regulates endoplasmic reticulum-mediated calcium signaling and cellular stress. Hum Mol Genet 19, 111–121, doi:10.1093/hmg/ddp471 (2010).

17 Carreras-Sureda, A. et al. ORMDL3 modulates store-operated calcium entry and lymphocyte activation. Hum Mol Genet 22, 519–530, doi:10.1093/hmg/dds450 (2013).

18 Moffatt, M. F. et al. A large-scale, consortium-based genomewide association study of asthma. N Engl J Med 363, 1211–1221, doi:10.1056/NEJMoa0906312 (2010).

19 Moffatt, M. F. et al. Genetic variants regulating ORMDL3 expression contribute to the risk of childhood asthma. Nature 448, 470–473, doi:10.1038/nature06014 (2007).

20 Bouzigon, E. et al. Effect of 17q21 variants and smoking exposure in early-onset asthma. N Engl J Med 359, 1985–1994, doi:10.1056/NEJMoa0806604 (2008).

21 Galanter, J. et al. ORMDL3 gene is associated with asthma in three ethnically diverse populations. Am J Respir Crit Care Med 177, 1194–1200, doi:10.1164/rccm.200711-1644OC (2008).

22 Barrett, J. C. et al. Genome-wide association study and meta-analysis find that over 40 loci affect risk of type 1 diabetes. Nat Genet 41, 703–707, doi:10.1038/ng.381 (2009).

23 McGovern, D. P. et al. Genome-wide association identifies multiple ulcerative colitis susceptibility loci. Nat Genet 42, 332–337, doi:10.1038/ng.549 (2010).

24 Liu, X. et al. Genome-wide meta-analyses identify three loci associated with primary biliary cirrhosis. Nat Genet 42, 658–660, doi:10.1038/ng.627 (2010).

25 Bugajev, V. et al. Negative regulatory roles of ORMDL3 in the FcepsilonRI-triggered expression of proinflammatory mediators and chemotactic response in murine mast cells. Cell Mol Life Sci 73, 1265–1285, doi:10.1007/s00018-015-2047-3 (2016).

26 Pelleymounter, M. A. et al. Effects of the obese gene product on body weight regulation in ob/ob mice. Science 269, 540–543 (1995).

27 Harris, R. B. et al. A leptin dose-response study in obese (ob/ob) and lean (+/?) mice. Endocrinology 139, 8–19, doi:10.1210/endo.139.1.5675 (1998).

28 Engin, F., Nguyen, T., Yermalovich, A. & Hotamisligil, G. S. Aberrant islet unfolded protein response in type 2 diabetes. Sci Rep 4, 4054, doi:10.1038/srep04054 (2014).

29 Engin, F. et al. Restoration of the unfolded protein response in pancreatic beta cells protects mice against type 1 diabetes. Sci Transl Med 5, 211ra156, doi:10.1126/scitranslmed.3006534 (2013).

30 Engin, F. ER stress and development of type 1 diabetes. J Investig Med 64, 2–6, doi:10.1097/JIM.0000000000000229 (2016).

31 Bonzon-Kulichenko, E. et al. Central leptin regulates total ceramide content and sterol regulatory element binding protein-1C proteolytic maturation in rat white adipose tissue. Endocrinology 150, 169–178, doi:10.1210/en.2008-0505 (2009).

32 Shimabukuro, M. et al. Lipoapoptosis in beta-cells of obese prediabetic fa/fa rats. Role of serine palmitoyltransferase overexpression. J Biol Chem 273, 32487–32490, doi:10.1074/jbc.273.49.32487 (1998).

33 Unger, R. H. & Roth, M. G. A new biology of diabetes revealed by leptin. Cell Metab 21, 15–20, doi:10.1016/j.cmet.2014.10.011 (2015).

34 Sloan, C. et al. Central leptin signaling is required to normalize myocardial fatty acid oxidation rates in caloric-restricted ob/ob mice. Diabetes 60, 1424–1434, doi:10.2337/db10-1106 (2011).

35 Roelants, F. M., Breslow, D. K., Muir, A., Weissman, J. S. & Thorner, J. Protein kinase Ypk1 phosphorylates regulatory proteins Orm1 and Orm2 to control sphingolipid homeostasis in Saccharomyces cerevisiae. Proc Natl Acad Sci U S A 108, 19222–19227, doi:10.1073/pnas.1116948108 (2011).

36 Sun, Y. et al. Orm protein phosphoregulation mediates transient sphingolipid biosynthesis response to heat stress via the Pkh-Ypk and Cdc55-PP2A pathways. Mol Biol Cell 23, 2388–2398, doi:10.1091/mbc.E12-03-0209 (2012).

37 Paulenda, T. & Draber, P. The role of ORMDL proteins, guardians of cellular sphingolipids, in asthma. Allergy 71, 918–930, doi:10.1111/all.12877 (2016).

38 Hagen-Euteneuer, N., Lutjohann, D., Park, H., Merrill, A. H., Jr. & van Echten-Deckert, G. Sphingosine 1-phosphate (S1P) lyase deficiency increases sphingolipid formation via recycling at the expense of de novo biosynthesis in neurons. J Biol Chem 287, 9128–9136, doi:10.1074/jbc.M111.302380 (2012).

39 Kiefer, K. et al. Coordinated regulation of the orosomucoid-like gene family expression controls de novo ceramide synthesis in mammalian cells. J Biol Chem 290, 2822–2830, doi:10.1074/jbc.M114.595116 (2015).

