## Supplementary figures and images for "Differential expression of Ormdl genes in the islets of mice and humans with obesity"

### Supplemental Figures

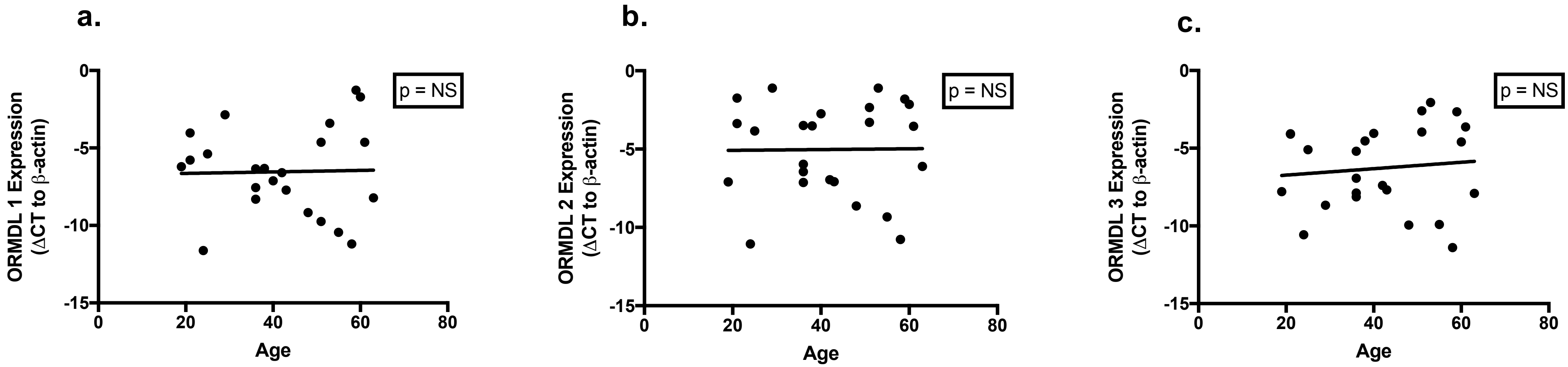

**Suppl. Fig2.**

**a.**

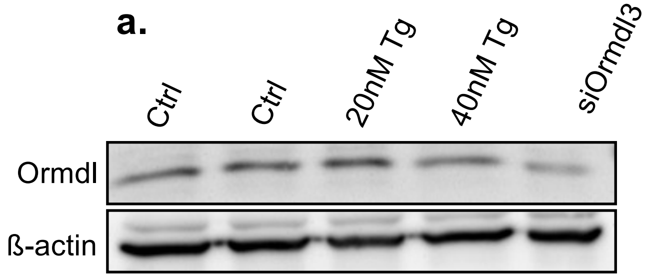


**b. Ormdl**

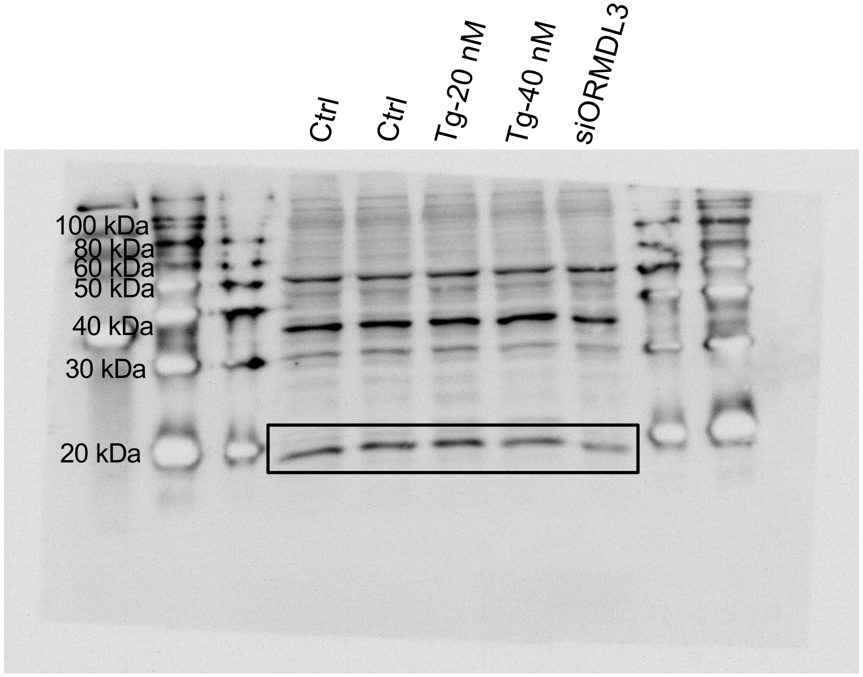


**c. B-actin**

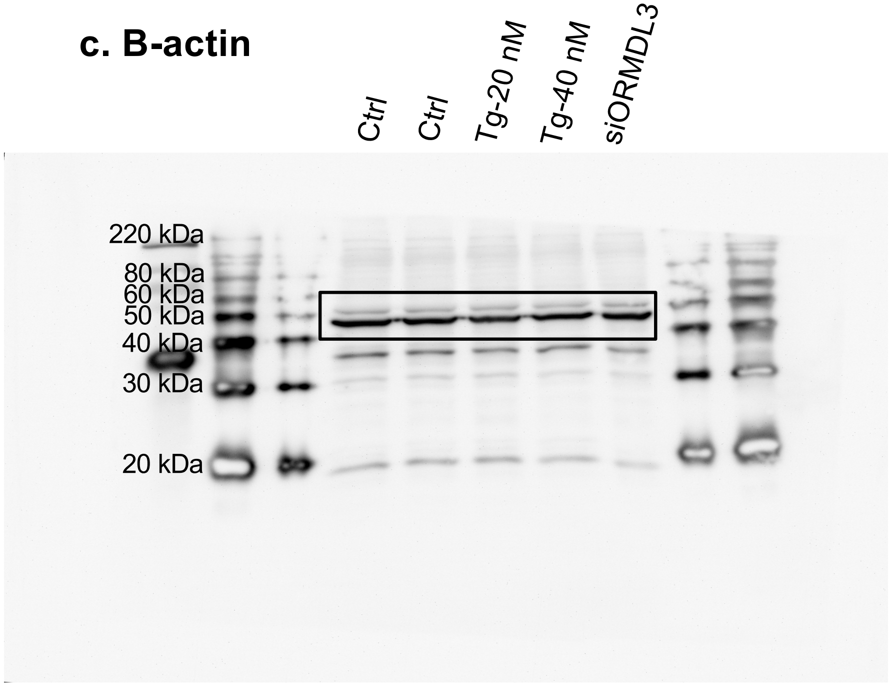
