## Supplementary material for "Differential expression of Ormdl genes in the islets of mice and humans with obesity": Supplemetal Figure Legends

**Supplementary information:**

Supplementary figure 1

Supplementary figure 2

**Supplemental Methods:**

**Cell culture**

INS-1 832/3 cells were cultured in RPMI 1640 supplemented with penicillin, streptomycin, 2 mM glutamine, 10mM HEPES, 1mM sodium pyruvate, 50 μM β-ME, and 10% FBS. The cells were maintained at 37°C in 5% $CO_2$ atmosphere and treated with 20-40 nM thapsigargin (Sigma-Aldrich), or with scrambled or siRNA against rat Ormld3 (Dharmacon).

**Western blot**

Cells were lysed in RIPA buffer (50 mM Tris, pH 7.4, 150 mM, NaCl, 5 mM EDTA, pH 8.0, 30 mM NaF, 1 mM Na3VO4, 40 mM β-glycerophosphate, 0.1 mM PMSF, protease inhibitors, 10% glycerol and 1% Nonidet-P40). Protein concentration of cellular lysates were determined using BCA Protein Assay Reagent (Pierce, Rockford, IL). Thirty-to-forty-five micrograms of protein were separated on a 5-12% Tris-acetate gel and electrophoretically transferred to PVDF membranes (Millipore, Billerica, MA). Membranes where then incubated with primary antibodies against sXBP1 (BioLegend), GRP78 (Cell Signaling Technology), Ormdl (TPF, gift of Dr. Petr Draber, Academy of Sciences of the Czech Republic), or β-actin (Cell Signaling Technology), followed by the appropriate secondary antibody.

### Supplementary Figure Legends

**Supplementary Figure 1. Human islet ORMDL expression is not correlated with donor age.** Scatter plots for **a.** ORMDL1, **b.** ORMDL2, **c.** ORMDL3 expression vs. age for all donors. Ns: not statistically significant.

**Supplementary Figure 2. Ormdl antibody specificity and expression in lean and obese mouse islets. a-c**. The specificity of a pan-Ormdl antibody was validated using the rat insulinoma cell line, INS-1 832/3, after transfection with siRNA against Ormdl3 (siOrmdl3) or scrambled control, as well as under stressed (thapsigargin treatment: Tg) or non-stressed conditions.
